# Co-phagocytosis of VEGFA with HER2-overexpressing cancer cells induced by HER2-VEGFA BsAb improves antitumor responses

**DOI:** 10.1101/2025.04.22.648615

**Authors:** Yang Lu, Songbo Qiu, Zhen Fan

## Abstract

We conceived of a new anti-tumor mechanism of action by which a soluble target in tumor microenvironment, such as a tumor-driving growth factor, can be phagocytized along with cancer cells via antibody-dependent cellular phagocytosis (ADCP) using an antibody bispecific for the soluble target and a solid target overexpressed on cancer cell surface. In this study, we explored the conception through engineering a pair of bispecific antibodies (BsAbs) co-targeting human epidermal growth factor receptor-2 (HER2) and human or mouse vascular endothelial growth factor A (VEGFA). We showed that the HER2-VEGFA BsAbs but not the parental antibodies alone or in combination induced co-phagocytosis of VEGFA and HER2-overexpressing cancer cells by tumor-associated macrophages via ADCP. In both immunocompromised and immunocompetent mice with aggressive tumors, the BsAbs demonstrated a greater anti-metastasis activity and had a greater survival benefit than the parental antibodies, alone or in combination, in a manner dependent on Fcγ receptors on the macrophages. Our results provide the proof-of-concept to induce VEGFA co-phagocytosis using HER2-VEGFA BsAbs to achieve enhanced antitumor activities by leveraging overexpressed HER2 on cancer cell surface. Our findings warrant clinical testing of the humanized BsAb to treat metastasis and recurrence of HER2-overexpressing solid tumors that respond to anti-VEGFA therapy.

**Graphical Abstract:** 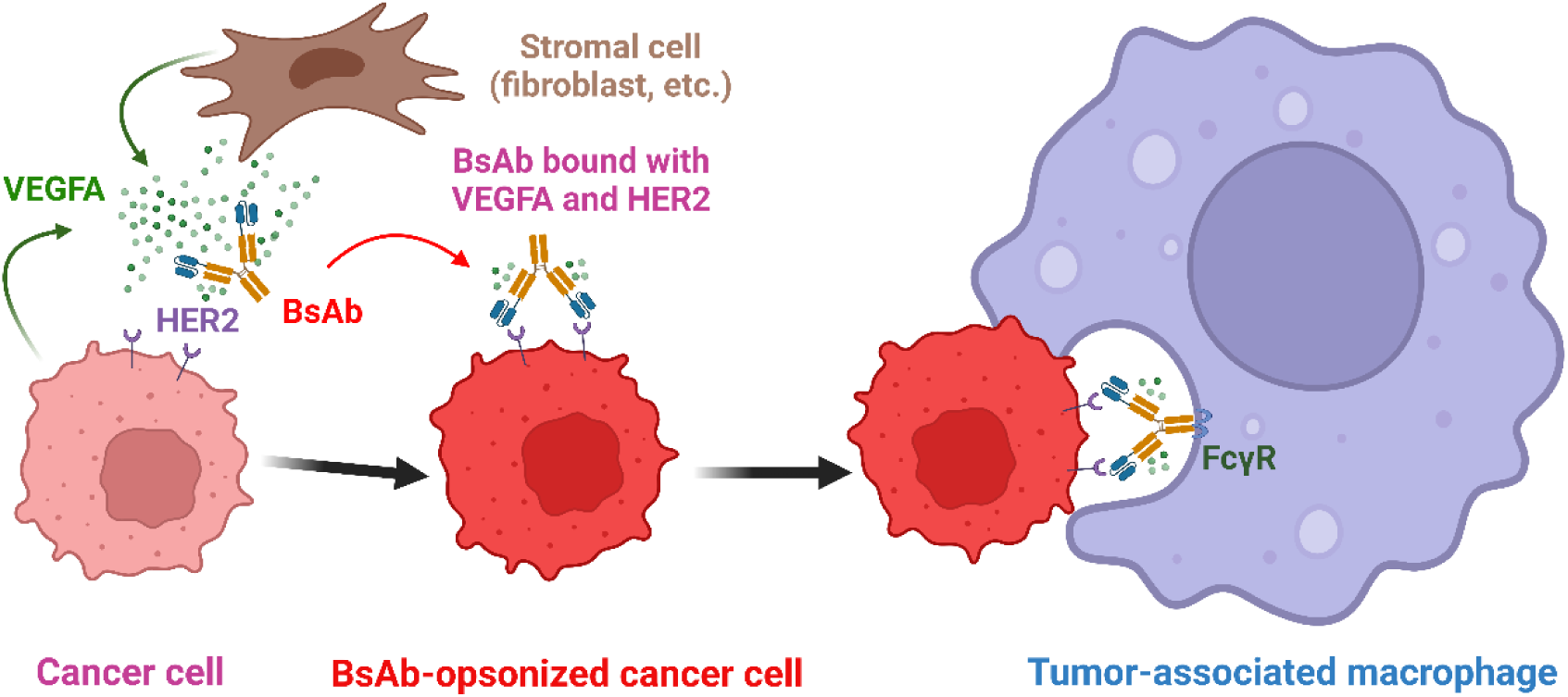

## Introduction

Bispecific antibodies (BsAbs) represent a new class of therapeutic antibodies that bind to and exert therapeutic activities against 2 different targets or 2 different epitopes on the same target (1, 2). Several bispecific targeting strategies for BsAbs have been explored (3, 4). The so-called biparatopic BsAbs bind to 2 non-overlapping epitopes on the same target, a strategy that can enhance binding avidity. Other BsAbs bind in a temporospatially coordinated manner to 2 different targets that function cooperatively, complementarily, or even redundantly, such as targets involved in immune checkpoint regulation, tumor heterogeneity, or ligand redundancy, and may thereby be more effective than simple combination of the 2 parental antibodies (4). Particularly compelling are BsAbs that exert new therapeutic activities through mechanisms that do not exist with simple combination of the 2 parental antibodies (4). Such BsAbs have been called *obligatory BsAbs* or *obligate BsAbs* because BsAb binding to 2 different antigen-binding sites is obligatory for gaining a new therapeutic mechanism of action (4). A successful example of such a BsAb is blinatumomab, a CD3- and CD19-binding BsAb that bridges CD3 on T cells to CD19 overexpressed on leukemia cells and thereby acts as a T-cell engager to direct the cytotoxic activity of T cells to the leukemia cells (5, 6). Another bispecific T-cell engager targeting CD3 and DLL3 was recently granted accelerated approval by US Food and Drug Administration (FDA) for treating DLL3-overexpressing small-cell lung cancer in 2024 (7). As of March 2025, 9 BsAbs are approved by the FDA, of which 7 are approved in oncology (8). More than 50 BsAbs have been investigated in clinical trials, and more than 180 BsAbs are in preclinical development (9).

Antibody-dependent cellular phagocytosis (ADCP) is a process wherein antibody-opsonized cells, pathogens, and sometimes soluble proteins are engulfed by macrophages following antibody’s Fc-mediated engagement of the Fc receptors on macrophages (10, 11). In this paper, we conceived of a new type of anti-tumor mechanism of action of BsAb designed to have one binding site for a solid target, such as a receptor or marker, overexpressed on cancer cell surface and the other binding site for a soluble target, such as a cytokine or other growth factor that is proven to be a key driver of tumor growth, metastasis, and/or immunosuppression in the tumor microenvironment (TME). We here coined the term *co-phagocytosis* to describe a process through which soluble targets in the TME are anchored to cancer cells by the new type of BsAb engineered as described, leading to internalization and degradation of the soluble targets in the macrophages via BsAb-induced ADCP of co-targeted cancer cells that overexpress a solid target on cell surface. We hypothesized that more soluble target molecules will be co-phagocytosed via BsAb-induced ADCP of cancer cells, because of overexpression of the solid target on the cancer cells, than via ADCP induced by an antibody specific for the soluble target only.

To validate the concept of co-phagocytosis through this bispecific targeting strategy, we selected human epidermal growth factor receptor-2 (HER2) and vascular endothelial growth factor A (VEGFA) because of proven clinical significance of these 2 targets and an existing pipeline of respective antibody drugs already clinically approved. HER2 was initially found to be overexpressed on breast cancer cells and is now emerging as a promising target for genomically informed therapy across a variety of solid tumor types (12, 13). VEGFA is well documented as a key cancer driver that is secreted by both cancer cells and cancer stromal cells in the TME and not only promotes tumor angiogenesis (14, 15) but also suppresses tumor immune responses (16, 17). In our studies, we generated and characterized HER2-VEGFA bispecific IgG1 to enable VEGFA co-phagocytosis through simultaneous binding to HER2 and VEGFA. We evaluated the therapeutic activities of HER2-VEGFA BsAbs against aggressive mouse tumors implanted in both immunocompromised nude mice and immunocompetent transgenic mice that are immunotolerant to HER2. In addition, we assessed the extent to which the therapeutic activity of HER2-VEGFA BsAbs depends on ADCP through co-administration of a Fcγ receptor (FcγR)-blocking antibody.

## Results

### Physicochemical properties and functional characterization of a BsAb platform

We opted for an scFv-IgG BsAb format, in which an scFv derived from a parental antibody targeting a cell surface marker is fused recombinantly to the N-terminus of the heavy chain of a parental antibody targeting a soluble molecule; this platform is referred to herein as heavy chain variable domain (VH)-modified-with-scFv (VHS) (Figure 1A). With the VHS platform, a trastuzumab-bevacizumab-VHS (TB-VHS) BsAb was engineered in a human IgG1 framework by fusing the scFv sequence of trastuzumab to the N-terminus of the VH for bevacizumab. As shown in Figure 1B, TB-VHS had a higher molecular weight than commercial trastuzumab or bevacizumab as a result of fusion of the trastuzumab-derived scFv to bevacizumab. The affinity of TB-VHS for HER2 was similar to the affinity of trastuzumab for HER2, and the affinity of TB-VHS for human VEGFA was similar to the affinity of bevacizumab for human VEGFA (Figure 1C). There was no noticeable loss of binding affinity to HER2 or to VEGFA by TB-VHS after incubation at 50°C for 1 h, compared to the binding affinity of TB-VHS kept at 4°C and to that of the respective parental antibodies treated in the same way (Figure 1D), indicating that the protein structure of the VHS platform was stable. With the VHS platform, we detected greater than 60% of the maximal binding of TB-VHS to VEGFA in the presence of an excess of HER2 extracellular domain (ECD) recombinant protein up to 10 times the concentration of VEGFA and vice versa (Figure 1E).

**Figure 1.**
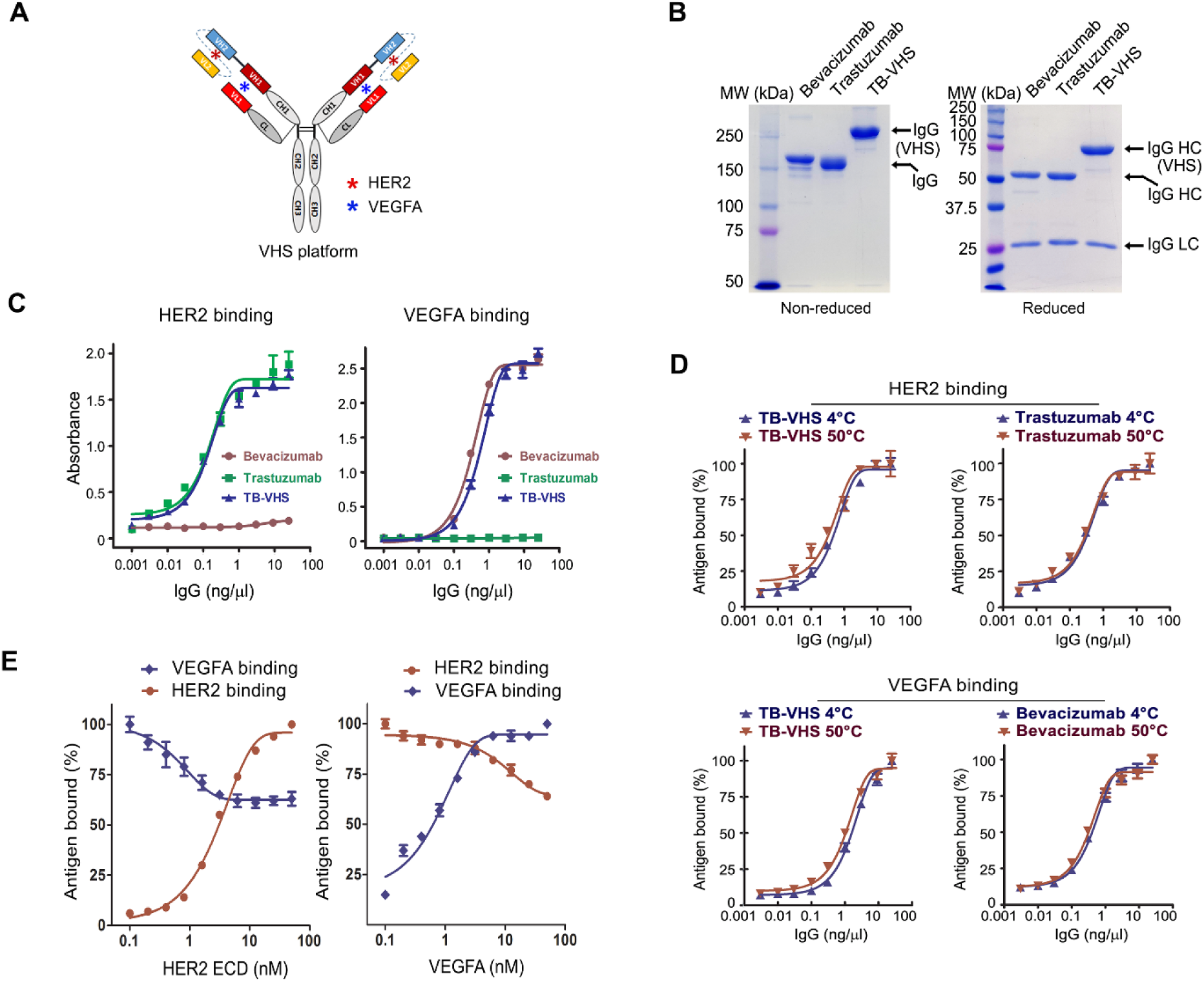
Physicochemical properties and functional characterization of TB-VHS. **(A)** Schematic illustration of BsAb in the VHS platform. The blue asterisks represent VEGFA to which the BsAb can bind, and the red asterisks represent HER2 to which the BsAb can bind. **(B)** Coomassie blue–stained gels of TB-VHS and its parental antibodies separated by SDS-PAGE under reducing (right) and nonreducing (left) conditions. HC, heavy chain; LC, light chain; MW, molecular weight. **(C)** Specific binding of TB-VHS and its parental antibodies to HER2 and human VEGFA detected by ELISA. For detecting HER2 binding, HER2 ECD recombinant protein–coated 96-well microplates were used to capture the antibodies, and antibodies were detected by HRP-labeled anti-human IgG antibody. For detecting VEGFA binding, rabbit anti-human Fc antibody–coated 96-well microplates were used to capture the antibodies, and antibodies were incubated with biotinylated human VEGFA and detected by streptavidin-HRP conjugate. **(D)** Thermal stability of TB-VHS. TB-VHS, bevacizumab, and trastuzumab were incubated in a water bath at 50°C for 1 hour, and then binding of these antibodies and antibodies stored at 4°C to HER2 and human VEGFA was detected by ELISA as in C. **(E)** Competitive binding of TB-VHS to HER2 and human VEGFA. TB-VHS (5 nM) was incubated with either 5-nM biotinylated VEGFA and increasing concentrations of HER2 ECD recombinant protein (left) or 5-nM HER2 ECD recombinant protein and increasing concentrations of VEGFA (biotinylated and unlabeled) (right) in a solution at 4°C for 1 hour. Then, separate 96-well microplates coated with rabbit anti-human Fc antibody were used to capture TB-VHS. Binding of VEGFA to TB-VHS was detected by streptavidin-HRP, and binding of HER2 ECD to TB-VHS was detected by a biotinylated anti-HER2 antibody and then streptavidin-HRP.

Next, we engineered trastuzumab-G6.31-VHS (TG-VHS) BsAb by fusing the scFv sequence of trastuzumab to the N-terminus of the VH for G6.31, a monoclonal antibody that targets both human and murine VEGFA (18, 19), in a mouse IgG2a framework to permit preclinical studies in immunocompetent mice. Figure 2 compares TB-VHS and TG-VHS for bispecific binding to HER2 and human or mouse VEGFA on live cells by flow cytometry analysis. Both TB-VHS and TG-VHS bound to HER2 on SKBR3 human breast cancer cells, detected respectively by an FITC-labeled anti-human IgG antibody and an FITC-labeled anti-mouse IgG (Figure 2, upper and lower left). Human VEGFA but not mouse VEGFA was detected on TB-VHS-treated SKBR3 cells (Figure 2, upper middle and right), because the anti-VEGFA component of TB-VHS is derived from bevacizumab, which binds only to human VEGFA (20, 21). By contrast, both human VEGFA and mouse VEGFA were detected on TG-VHS-treated cells (Figure 2, lower middle and right). Together, these results confirm that TB-VHS is a humanized antibody bispecific for HER2 and human VEGFA but not mouse VEGFA, whereas TG-VHS is a mouse antibody bispecific not only for HER2 and human VEGFA but also for HER2 and mouse VEGFA.

**Figure 2.**
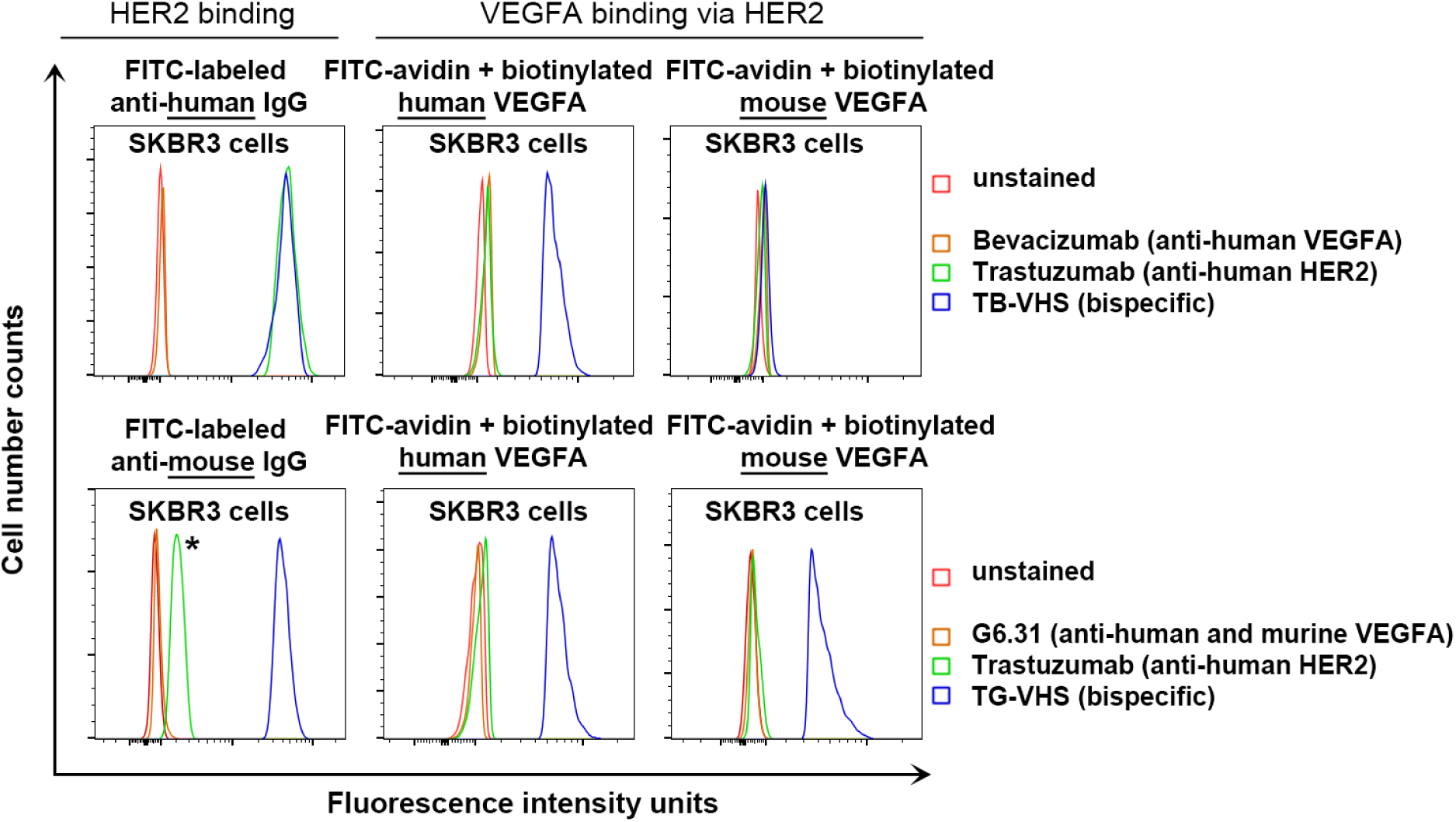
Detection of bispecific binding of TB-VHS and TG-VHS to human and mouse VEGFA and to HER2 on live cells by flow cytometry analysis. After incubation at 4°C for 1 hour with TB-VHS (upper panel) or TG-VHS (lower panel) or respective parental antibodies as shown, HER2-overexpressing SKBR3 cells were incubated with FITC-labeled anti-human IgG antibody (upper panel, left column) or FITC-labeled anti-mouse IgG antibody (lower panel, left column); FITC-avidin plus biotinylated human VEGFA (middle column); or FITC-avidin plus biotinylated mouse VEGFA (right column). The asterisk indicates minor cross-reaction of the FITC-labeled anti-mouse IgG antibody with trastuzumab (humanized IgG).

### Co-phagocytosis of fibroblast-secreted VEGFA with HER2-overexpressing cancer cells via HER2-VEGFA BsAb-induced ADCP

To demonstrate our proposed mechanism of VEGFA co-phagocytosis via ADCP of HER2-overexpressing cells by the BsAb, we co-cultured 4T1 mouse mammary tumor cells lentivirally transduced to express HER2, firefly luciferase, and mCherry protein (termed 4T1-HLC cells) with RAW264.7 mouse macrophages in fresh culture medium supplemented with conditioned medium from culture of NIH3T3 fibroblasts lentivirally transduced to secrete human VEGFA fused with enhanced green fluorescent protein (EGFP) (termed NIH3T3-VG cells) or from culture of the fibroblasts lentivirally transduced with the EGFP construct only (termed NIH3T3-G cells). Flow cytometry was used to detect mCherry and EGFP fluorescent signals in the RAW264.7 macrophages following each of 5 treatments: 1) control mouse IgG; 2) G6.31.2a, constructed on the basis of G6.31 sequences but with its Fc domain re-engineered in the mouse IgG2a framework; 3) 4D5.2a, constructed on the basis of trastuzumab but with its Fc domain re-engineered in the mouse IgG2a framework; 4) G6.31.2a plus 4D5.2a; or 5) TG-VHS. Figure 3A (upper panel) shows a basal level of phagocytosis or levels of ADCP of 4T1-HLC cells (mCherry-positive) by RAW264.7 macrophages (F4/80-APC-positive) after each of the 5 treatments in the co-cultures supplemented with the control medium (left panel) or VEGFA medium (right panel). A basal level of phagocytosis was detected following control mouse IgG treatment of the co-cultures supplemented with the control medium or VEGFA medium (7% and 5%, respectively). A similar level of phagocytosis was observed upon G6.31.2a treatment of the co-cultures supplemented with the control medium or VEGFA medium (8% and 6%, respectively), but the level of phagocytosis was significantly increased upon treatments that included a HER2-targeting antibody, including 4D5.2a (20% and 19%, respectively), G6.31.2a plus 4D5.2a (19% and 17%, respectively), and TG-VHS (24% and 21%, respectively), indicating increased levels of phagocytosis of 4T1-HLC cells by RAW264.7 macrophages via ADCP. Figure 3A (lower panel) shows levels of co-phagocytosis of VEGFA-EGFP with 4T1-HLC cells by RAW264.7 macrophages under the same treatment conditions as in the upper panel of Figure 3A. Compared with the other treatments, TG-VHS treatment led to a significant increase in the subpopulation of cells dual positive for F4/80 and VEGFA-EGFP (Q2), from < 5% with the other treatments to 14% with TG-VHS (Figure 3A, right top row in the lower panel). Further gating of Q2 showed that 79% of the VEGFA-EGFP-positive macrophages were also positive for mCherry (Figure 3A, right middle row in the lower panel), confirming that the majority of VEGFA-EGFP in the macrophages following TG-VHS treatment was co-phagocytosed with 4T1-HLC cells (mCherry-positive). Further gating of the cells positive for VEGFA-EGFP but negative for F4/80 (Q1) confirmed that 95% of these cells were also mCherry-positive (Figure 3A, right bottom row in the lower panel), indicating that these cells were 4T1-HLC cells bound to VEGFA-EGFP but yet to be phagocytosed by RAW264.7 macrophages. Figure 3B shows representative confocal imaging of 4T1-HLC cells and RAW246.7 cells co-cultured in the presence of VEGFA-EGFP collected from the conditioned medium of NIH3T3-VG cell culture as described in Figure 3A. The VEGFA-EGFP, which is a soluble protein undetectable in the co-culture treated with control mouse IgG, became detectable after it was anchored on 4T1-HLC cell surface upon the BsAb treatment and was subsequently co-phagocytosed along with 4T1-HLC cells by RAW264.7 cells via BsAb-induced ADCP. Together, these results support a working model depicted in Figure 3C.

**Figure 3.**
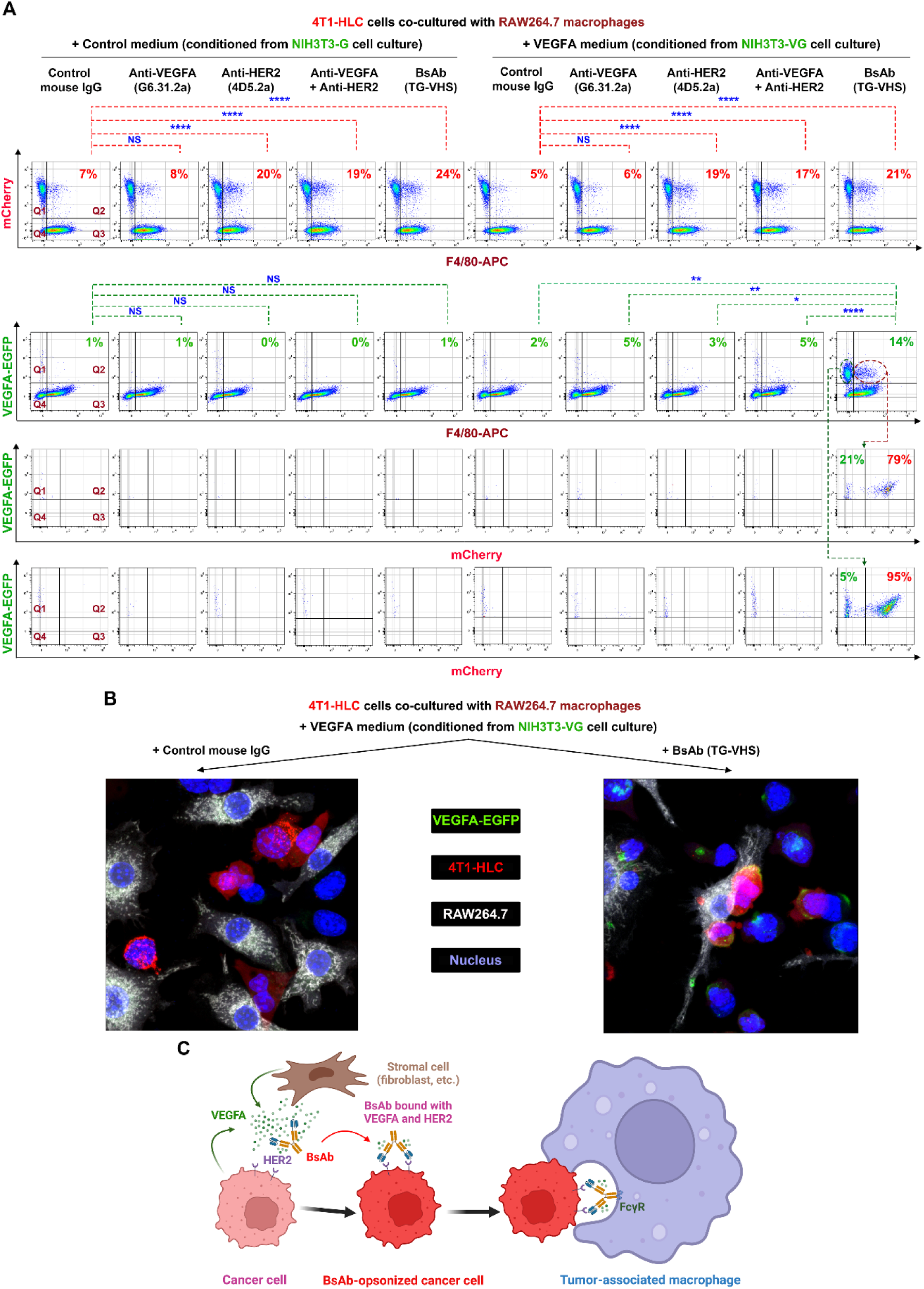
Co-phagocytosis of fibroblast-secreted VEGFA with HER2-overexpressing cancer cells via HER2-VEGFA BsAb-induced ADCP in co-culture with macrophages. **(A)** 4T1-HLC cells and RAW264.7 macrophages (at a 1:5 ratio of cell number) were co-cultured in fresh medium supplemented with conditioned medium (at a 1:1 ratio of volume) from culture of NIH3T3-G cells or from culture of NIH3T3-VG cells and were treated with one of the antibodies (0.5µg/mL each) as indicated for 2 hours in a 37°C incubator with 5% CO2. The cells were then harvested and stained with APC-conjugated anti-F4/80 antibody for flow cytometry analysis. *Upper panel:* the co-cultured cells were gated for mCherry positivity in F4/80+ cells (indicating ADCP of HER2-overexpressing cells). The cells in Q2 indicate 4T1-HLC cells phagocytized by the macrophages in the co-culture. Phagocytosis % = Q2/[Q1+Q2]. *Lower panel:* in the top row, the co-cultured were gated for GFP positivity in F4/80+ cells (indicating VEGFA co-phagocytosis via ADCP of HER2-overexpressing cells). Co-phagocytosis % = Q2/[Q1+Q2]. In the middle row, GFP and F4/80 double-positive cell population was further gated for mCherry positivity. In the bottom row: The GFP-positive but F4/80-negative cell population was further gated for mCherry positivity. **(B)** RAW246.7 cells were stained with CellTrace Far Red fluorescent dye (shown in white color) and then seeded on a poly-D-lysine-coated cell culture dish (FluoroDish) overnight. Next day, 4T1-HLC cells (shown in red color) were added to the RAW246.7 cell culture and incubated for 3 hours in the presence of TG-VHS or control antibody as indicated, along with concentrated conditioned medium from NIH3T3-VG cell culture containing VEGFA-GEP (shown in green color). The cells were counterstained with DAPI fluorescent dye (shown in blue color) and then fixed with 2% polyformaldehyde prior to confocal cell imaging analysis. **(C)** A model depicting the process of VEGFA co-phagocytosis induced by the HER2-VEGFA BsAb (created with BioRender).

### In vivo activity of HER2-VEGFA BsAb against metastasis of 4T1-HL cells co-implanted with VEGFA-secreting fibroblasts in immunocompromised mice

We next asked if our proposed mechanism of action by the BsAb leads to an enhanced antitumor activity in mice co-implanted in the mammary fat pad with HER2-overexpressing 4T1-HL cells (similar to 4T1-HLC cells but without mCherry) with VEGFA-secreting NIH3T3-V fibroblasts (similar to NIH3T3-VG cells but secreting VEGFA rather than VEGFA-EGFP fusion protein). Because HER2 and human VEGFA are immunogenic in normal mice, we set up the co-implantation experiment in nude mice, which, despite being immunodeficient, possess functional macrophages required to mediate ADCP (22) and which also permit use of the humanized TB-VHS BsAb we developed in this study. When the tumors were detected by IVIS imaging, on day 4 after cell implantation, treatment was initiated with control human IgG, bevacizumab, trastuzumab, trastuzumab plus bevacizumab, or TB-VHS, with antibodies administered twice a week in line with practices commonly reported in the literature (23–26). To achieve a molar mass of TB-VHS equal to that of the parental mAbs at 100 μg/mouse when used alone and at 2×100 μg/mouse when used in combination, we opted for 150 μg of BsAb per mouse to account for the higher molecular weight (∼225 kDa) of TB-VHS than of the parental antibodies (∼150 kDa each) owing to the scFv fusion (Figure 1, A and B).

Tumor metastasis was detected on day 10, and shortly after, the mice started to die of massive metastasis. Simple combination of trastuzumab and bevacizumab produced no advantage over either single treatment alone in controlling the extent of metastasis; however, TB-VHS-treated mice had remarkably less extensive metastasis than mice in the other groups had on day 24 (Figure 4A). The mice in all groups were treated as scheduled until all mice in the control group had died and the number of mice remaining in any of the treatment groups became too small to permit statistical comparisons. On day 34, when all mice in the other groups had died, 4 of the 8 mice treated with TB-VHS remained alive (Figure 4B). However, tumors persisted in the surviving mice and, owing to the aggressiveness of the tumor model, the mice eventually died. Nevertheless, the results showed greater antitumor activity of TB-VHS than of the parental antibodies used alone or in combination.

**Figure 4.**
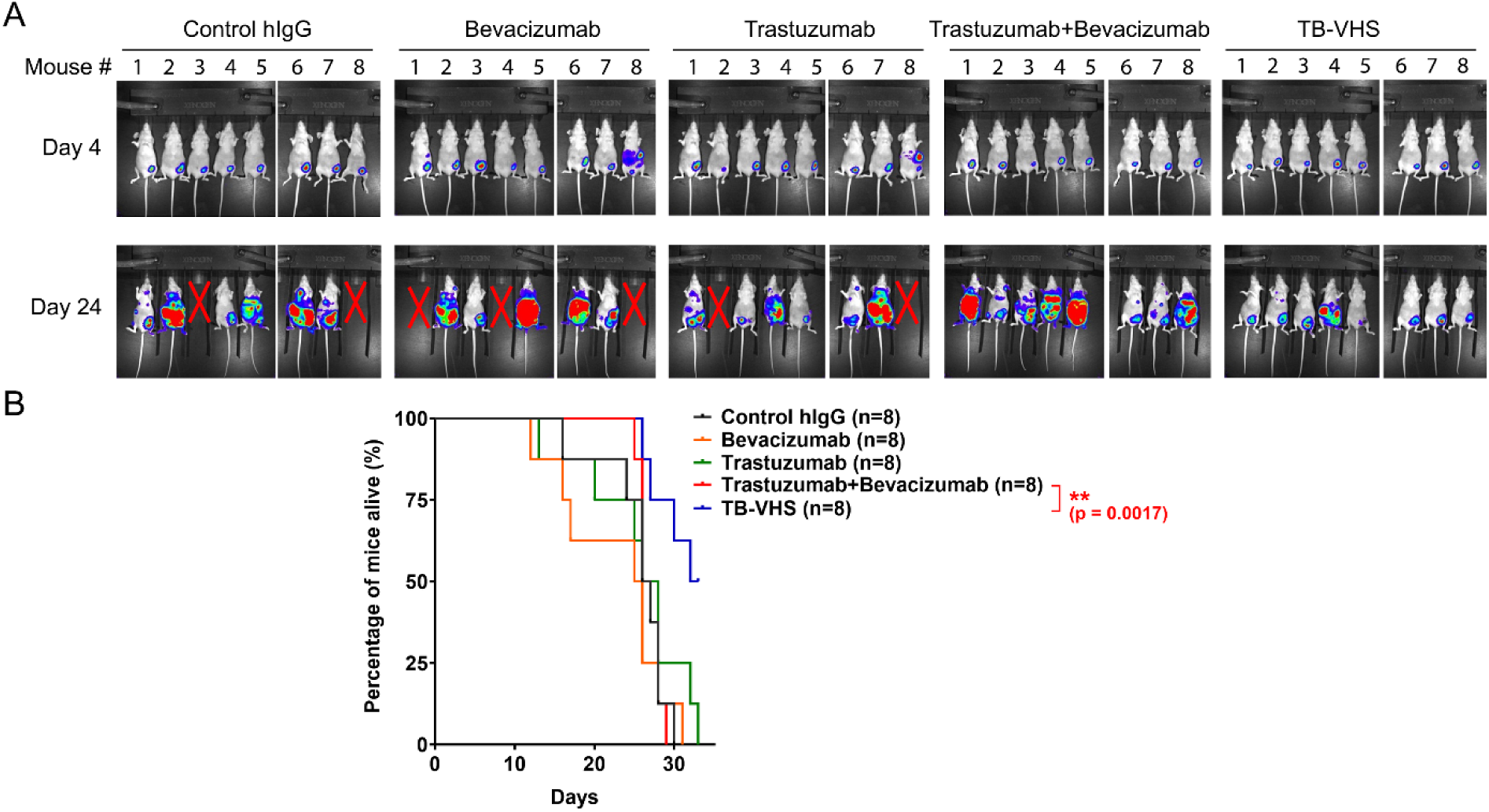
Anti-metastasis activity of TB-VHS against 4T1-HL tumors co-implanted with human VEGFA-secreting mouse fibroblasts in nude mice. **(A)** 4T1-HL mouse mammary tumor cells were co-implanted along with NIH3T3 fibroblasts transduced to express human VEGFA into the mammary fat pad of nude mice. The mice were divided into 5 groups with 8 mice per group. Starting on day 4, mice were injected intraperitoneally twice a week with control human IgG (hIgG) (100 μg/mouse), bevacizumab (100 μg/mouse), trastuzumab (100 μg/mouse), trastuzumab plus bevacizumab (100 μg/mouse + 100 μg/mouse), or TB-VHS (150 μg/mouse, equivalent to 100 μg in molar mass of conventional antibody). The mice were subjected to IVIS imaging on day 4 and day 24. *X* indicates mice that died before day 24. **(B)** Survival curves of the mouse treatment groups described in A.

### HER2-VEGFA BsAb-induced co-phagocytosis of recombinant VEGFA via ADCP of HER2-overexpressing cells co-cultured with primary bone marrow-derived macrophages

To further validate our working model, we harvested primary bone marrow cells from normal mice and subjected the cells to differentiation into bone marrow-derived macrophages (BMDM) by culture of the cells with macrophage colony-stimulating factor (M-CSF) for 7 days. The BMDM cells were left unpolarized or further polarized to the M1 type macrophages by treatment with a combination of interferon gamma (IFNγ) and lipopolysaccharide (LPS) or to the M2 type macrophages by treatment with a combination of interleukin (IL)-4, IL-10, and IL-13. Flow cytometry analysis of the BMDM cells shows a marked increase in the level of CD86 (a marker of M1 type macrophages) or CD206 (a marker of M2 type macrophages), confirming successful polarization of the BMDM cells after respective treatment (Figure 5A). The level of FcγRs detected by 2.4G2 antibody, which recognizes both FcγRIII/CD16 and FcγRII/CD32 (27, 28), was also found to be higher in the polarized BMDM cells than in the unpolarized cells. These unpolarized or M1- or M2-polarized BMDM cells were then co-cultured with syngeneic D5 mouse melanoma cells transduced to overexpress HER2 (D5-HER2) for further characterization of co-phagocytosis of a recombinant VEGFA (labeled with pH-sensitive pHrodo green) after overnight treatment with TG-VHS, combination of anti-VEGFA (G6.31.2a) and anti-HER2 (4D5.2a) antibodies or a control antibody. Unlike the VEGFA-EGFP protein in Figure 3, the pH-sensitive VEGFA green fluorescence was detectable by flow cytometry only when the VEGFA was internalized and transported to the endosomes or was phagocytosed and transported to the phagosomes in the treated cells. As shown in the top row in Figure 5B, a basal level of pH-sensitive VEGFA green fluorescence was detected in the BMDM cells (F4/80-positive) cultured alone, suggesting the presence of VEGFA receptors on the surface of the BMDM cells. An increase in the level of the VEGFA green fluorescence was detected in the BMDM cells co-cultured with D5-HER2 tumor cells (F4/80-negative) after treatment with the combination of the anti-VEGFA and anti-HER2 antibodies, which indicates additional VEGFA molecules opsonized by the anti-VEGFA antibody were endocytosed and transported to the phagosomes in the BMDM cells. Importantly, only in TG-VHS-treated co-culture was the VEGFA green fluorescence detected in both D5-HER2 tumor cells and the BMDM cells, and the level of VEGFA green fluorescence detected in the BMDM cells was higher after treatment with the BsAb than with the antibody combination. Similar findings but at higher levels were found in the co-culture of D5-HER2 tumor cells with the polarized BMDM cells, particularly with the M2 type BMDM cells, than with the unpolarized BMDM cells (Figure 5B, middle and bottom rows). These findings are consistent with the observation of increased levels of FcγRs in the M1- and M2-polarized BMDM cells (Figure 5A), which are key mediators in ADCP induction (29, 30). Together, these results further validate our working model illustrated in Figure 5C that TG-VHS BsAb opsonizes VEGFA and induces VEGFA co-phagocytosis by leveraging overexpressed HER2 on cancer cell surface via ADCP that can be mediated by both unpolarized and M1- or M2-polorized BMDM cells and that this mechanism of VEGFA co-phagocytosis is not actionable by the parental antibodies alone or in simple combination.

**Figure 5.**
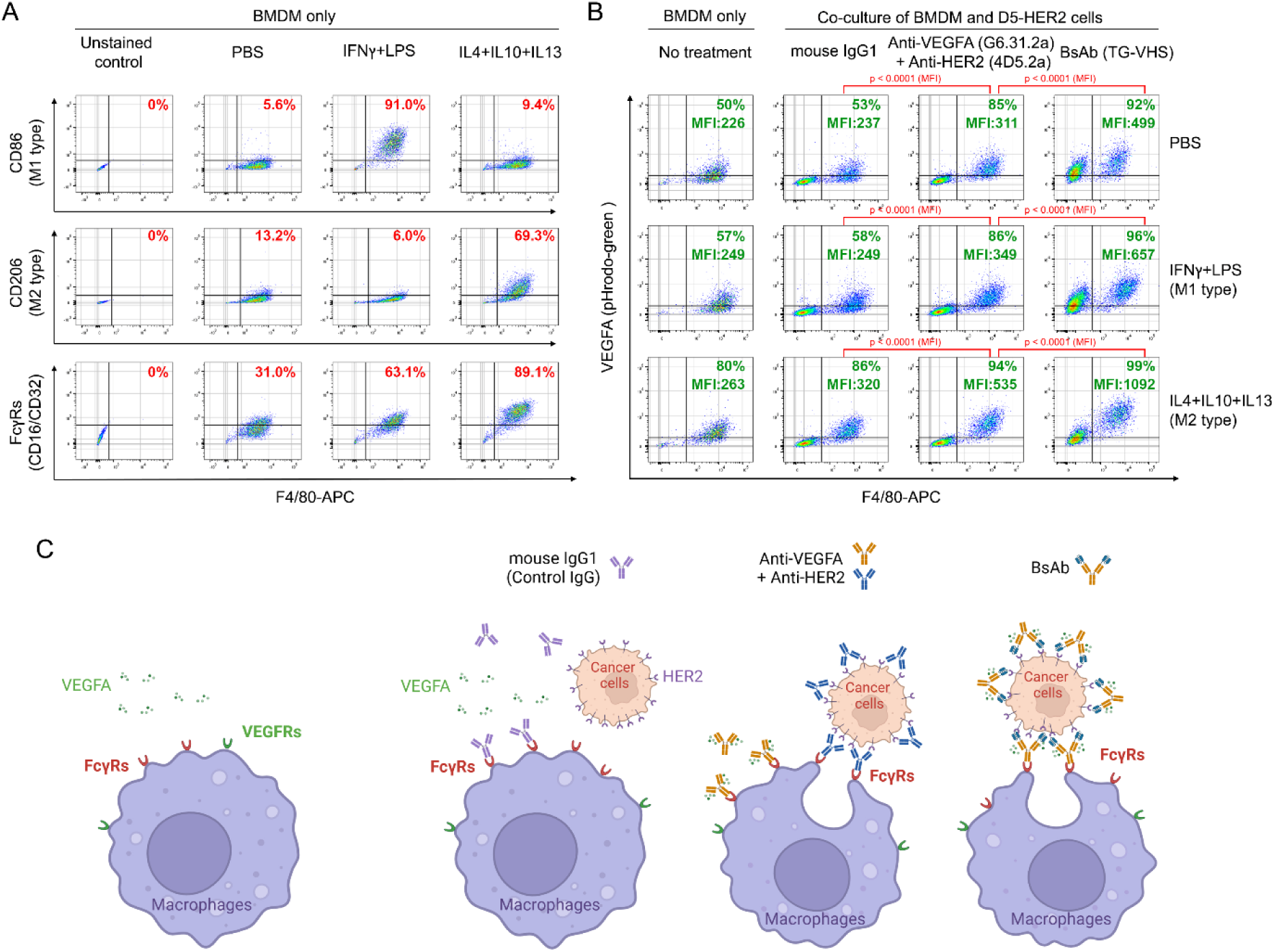
Analysis of VEGFA co-phagocytosis with HER2-overexpressing cancer cells via HER2-VEGFA BsAb-induced ADCP in co-culture with unpolarized and polarized BMDM cells. **(A)** BMDM cells were subjected to flow cytometry analysis for the levels of CD86 (an M1 marker) or CD206 (an M2 marker) on the cells after combination treatment with IFNγ (50 ng/mL) and LPS (1:1000 dilution of the stock) for 12 hours or with IL-4, IL-10, and IL-13 (50 ng/mL each) for 24 hours in culture as indicated. **(B)** BMDM cells unpolarized or polarized as shown in (A) were mixed with D5-HER2 cells (at a 1:2 ratio of cell number) in the presence of mouse IgG control antibody, combination of G6.31.2a and 4D5.2a, or TG-VHS (0.5µg/mL each). The cell mixture or the BMDM cells alone were incubated with a pHrodo green-labeled recombinant VEGFA (0.5µg/sample) for overnight. The cell samples were then subjected to flow cytometry analysis after staining with APC-conjugated anti-F4/80 antibody. The flow cytometry data were gated using FlowJo software package (v10) for VEGFA green positivity in F4/80-positive cells (BMDM) and F4/80-negative cells (D5-HER2). The gating strategies of flow cytometry analysis in (A) and (B) are shown in supplemental figures. **(C)** Models depicting the processes of phagocytosis and co-phagocytosis of VEGFA opsonized upon the treatments (created with BioRender).

### Improvement of tumor-free survival by HER2-VEGFA BsAb in immunocompetent hmHER2Tg mice with aggressive D5-HER2 tumors

To further investigate the extent to which the proposed mechanism of action by the BsAb leads to an enhanced antitumor activity, we implanted D5-HER2 syngeneic tumor cells in hmHER2Tg mice, a transgenic mouse line in C57BL/6 background that is immunocompetent but immunotolerant to HER2 (31–33). D5-HER2 tumor cells implanted subcutaneously in hmHER2Tg mice grow very aggressively that can kill the mice (similar to the killing of mice by 4T1-HL cells in Figure 4). In experiments similar to the one described in Figure 4 but with mouse antibodies used instead, the hmHER2Tg mice were randomly divided into 5 groups for treatment with control mouse IgG, G6.31.2a anti-VEGFA antibody, 4D5.2a anti-HER2 antibody, G6.31.2a plus 4D5.2a, or TG-VHS twice a week for 6 weeks, with treatment started on the day after D5-HER2 tumor cell implantation (Figure 6). Figure 6A shows tumor development in each mouse in each treatment group (n=15 in each group) from day 0 after tumor cell implantation. Tumor measurement was stopped after day 23, when the mice in the control group started dying. Within 52 to 122 days after tumor cell implantation, all 15 mice treated with the control mouse IgG had died (median survival, 28 days); furthermore, only 2 of 15 mice (13.3%) treated with G6.31.2a (median survival, 25 days), 4 of 15 mice (26.7%) treated with 4D5.2a (median survival, 39 days), and 5 of 15 mice (33.3%) treated with G6.31.2a plus 4D5.2a (median survival, 71 days) were tumor free at day 257 (Figure 6B). By contrast, 8 of 15 mice (53.3%) treated with TG-VHS were tumor-free (median survival, >257 days). However, despite the clear survival benefit of TG-VHS treatment, the difference in survival between the TG-VHS group and the G6.31.2a plus 4D5.2a group was not statistically significant, which prompted us to conduct another experiment with a re-designed treatment plan shown below in Figure 7.

**Figure 6.**
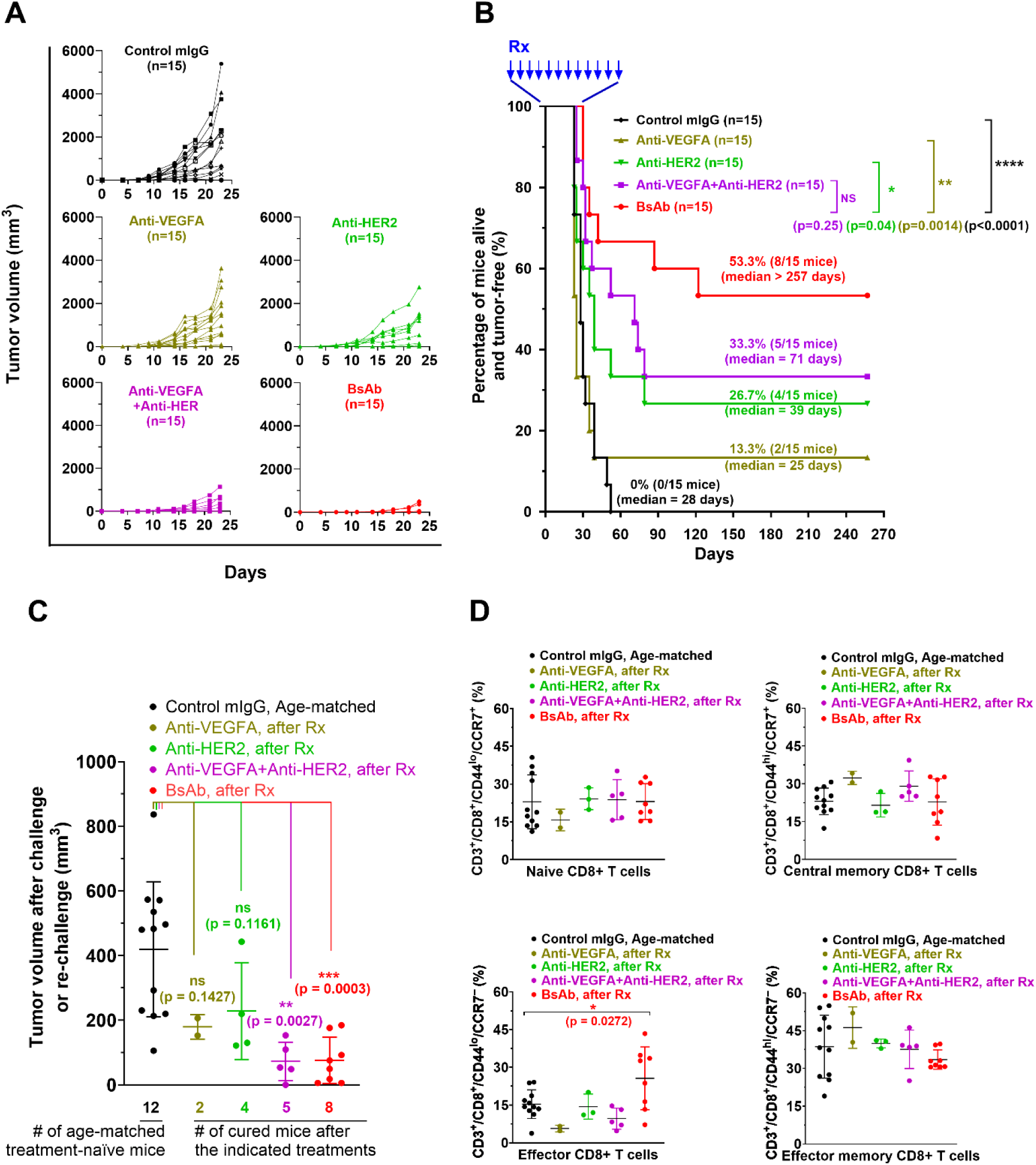
Improvement of tumor-free survival of hmHER2Tg mice with D5-HER2 tumors by TG-VHS compared with combination of anti-VEGFA and anti-HER2 antibodies. **(A)** Tumor growth in individual mice in each treatment group. D5-HER2 syngeneic tumor cells (2.5×10^5^ cells/mouse) were implanted subcutaneously into hmHER2Tg mice. The next day (day 1), the mice were randomly divided into 5 groups, and treatment was initiated. Mice received a control mouse IgG (mIgG) (100 µg/mouse), G6.31.2a (100 µg/mouse), 4D5.2a (100 µg/mouse), G6.31.2a plus 4D5.2a (100 µg/mouse+100 µg/mouse), or TG-VHS (150 µg/mouse, equivalent to 100 µg in molar mass of conventional antibody) intraperitoneally twice a week for 6 weeks. Tumor measurement was stopped after day 23, when mice started to die or had to be euthanized owing to moribund status. **(B)** Survival curves of the mouse treatment groups described in A. Treatments were stopped after 6 weeks, and the mice were monitored for survival up to 257 days. Rx, treatment. **(C)** Tumor challenge and rechallenge in hmHER2Tg mice. Age-matched treatment-naïve hmHER2Tg mice were challenged and the mice in A and B that remained tumor-free during the extended period (257 days) were rechallenged with equal amounts (2.5×10^5^ cells/mouse) of the same parental D5 tumor cells (without HER2 overexpression). Tumor growth was measured twice a week for 2 weeks. Plots show tumor volume in individual mice on day 14. Rx, treatment. **(D)** Immunophenotypic analysis of the spleen specimens from the mice in each group in C. Single cell suspensions were prepared for flow cytometry analysis for the relevant markers shown. Rx, treatment.

**Figure 7.**
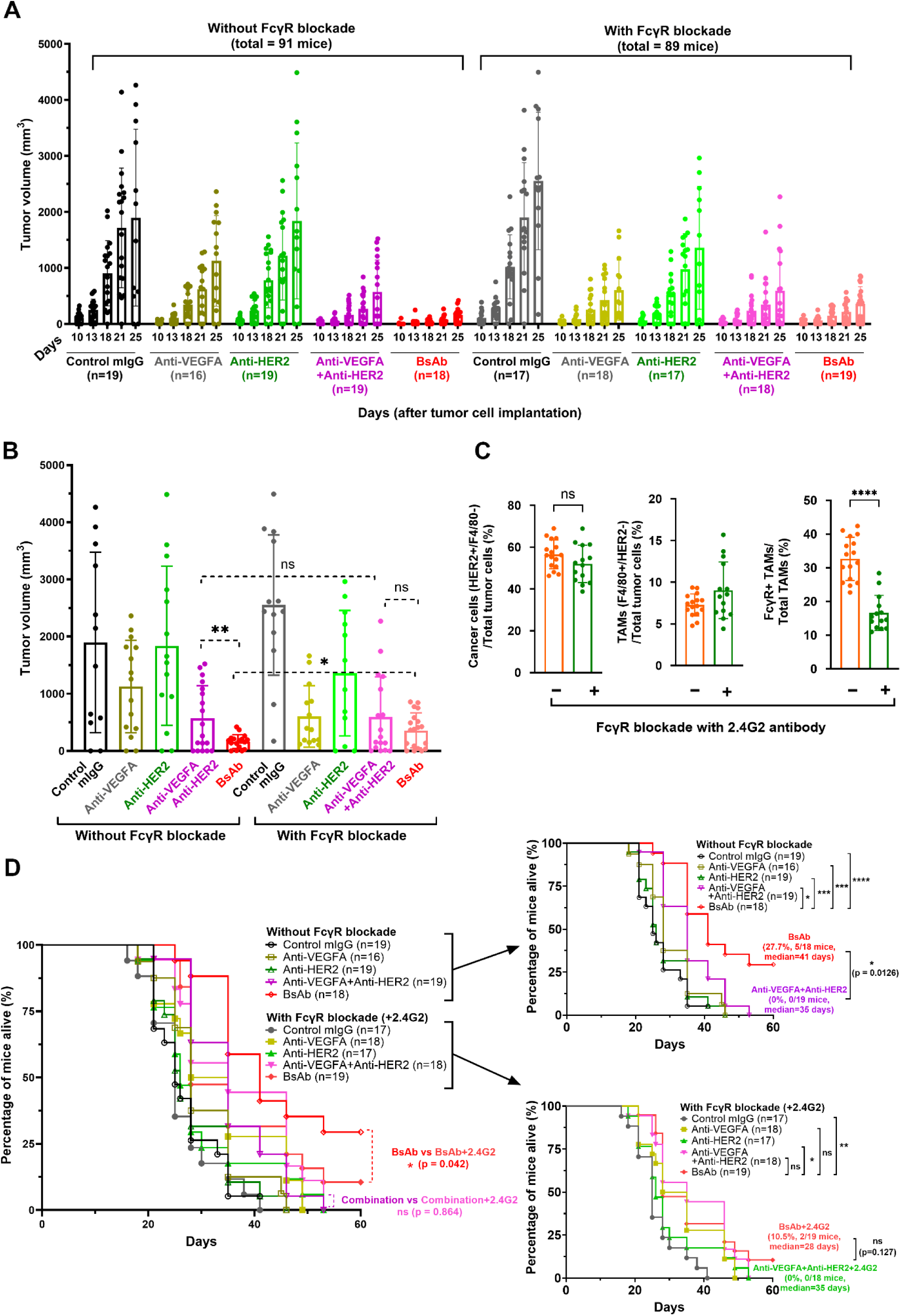
Dependence on FcγRs on TAMs in mediating an enhanced antitumor activity and survival benefit by TG-VHS compared with combination of anti-VEGFA and anti-HER2 antibodies in hmHER2Tg mice with D5-HER2 tumors. **(A)** Tumor growth in individual mice after the indicated treatments without and with FcγR blockade. D5-HER2 syngeneic tumor cells (2.5×10^5^ cells/mouse) were transplanted subcutaneously into hmHER2Tg mice. The mice were then randomly divided into 2 cohorts receiving 2.4G2 FcγR-blocking antibody (100 µg/mouse) or not on the same day. On day 3, the mice in each cohort were further randomly divided into 5 groups, and treatment was initiated. Mice received a control mouse IgG (mIgG) (100 µg/mouse), G6.31.2a anti-VEGFA antibody (100 µg/mouse), 4D5.2a anti-HER2 antibody (100 µg/mouse), G6.31.2a plus 4D5.2a (100 µg/mouse+100 µg/mouse), or TG-VHS (150 µg/mouse, equivalent to 100 µg in molar mass of conventional antibody) intraperitoneally twice a week for 5 weeks. Tumor measurement was stopped after day 25, when mice started to die or had to be euthanized owing to moribund status. **(B)** Statistical comparison of tumor growth on day 25 between the major groups of interest. **(C)** Analysis of result of pharmacological blockade of FcγR ex vivo. Shown are flow cytometry data from tumor samples collected from moribund mice that were euthanized on day 33. The samples within each cohort were grouped, and the grouped samples were subjected to flow cytometry analysis after staining with a mixture of antibodies including BV421-conjugated anti-HER2 antibody, APC-conjugated anti-F4/80 antibody, and PE-conjugated nonspecific goat-anti-rabbit IgG antibody. **(D)** Survival curves of all the mouse treatment groups described in A and B (left panel) and of the mouse treatment groups without (right panel, upper) and with (right panel, lower) FcγR blockade. ns, not significant. * p<0.05. **p<0.01, ****p<0.0001

Meanwhile, the mice in the various treatment groups in Figure 6A and 6B that were alive with tumor-free at the end of the experiment (day 257) were considered cured and were then re-challenged with equal amounts of parental D5 tumor cells without HER2. Age-matched treatment-naïve mice implanted with equal amounts of D5 tumor cells were used as controls. Compared with the growth of D5 tumor implants in the control mice, growth of D5 tumor implants in the cured mice treated with an anti-VEGFA activity (G6.31.2a alone, G6.31.2a plus 4D5.2a, or TG-VHS) was remarkably slower (Figure 6C), and the difference was statistically significant in the TG-VHS and combination treatment groups but not in the G6.31.2a alone group owing to the low number of mice that survived. All mice were euthanized 2 weeks after the rechallenge, and the spleens were collected for CD8^+^ T cell immunophenotype analysis. No significant differences were found in the subpopulations of naïve CD8^+^ T cells (CD3^+^/CD8^+^/CD44^lo^/CCR7^+^), central memory CD8^+^ T cells (CD3^+^/CD8^+^/CD44^hi^/CCR7^+^) or effector memory CD8^+^ T cells (CD3^+^/CD8^+^/CD44^hi^/CCR7^-^); however, the subpopulation of effector CD8^+^ T cells (CD3^+^/CD8^+^/CD44^lo^/CCR7^-^) was significantly more abundant in the mice treated with TG-VHS than in the mice in the other groups (Figure 6D), indicating that effector CD8^+^ T cells might have been activated in response to TG-VHS therapy.

### Dependence of HER2-VEGFA BsAb on FcγRs on the macrophages in mediating the antitumor activity and the survival benefit of the BsAb

To confirm the role of VEGFA co-phagocytosis via BsAb-induced ADCP (Figures 3 and 5) in improving antitumor responses to the BsAbs leading to a survival benefit (Figures 4 and 6), we examined the extent to which inhibition of ADCP affected tumor responses to the antibody treatments by treating the mice with and without co-administration with the 2.4G2 antibody to pharmacologically block FcγRs on tumor-associated macrophages (TAMs) in vivo. Following implantation of D5-HER2 tumor cells in hmHER2Tg mice, the mice were randomly assigned to 2 large cohorts, one receiving 2.4G2 antibody starting on day 0 and the other not receiving 2.4G2 antibody. Treatments were started on day 3, when tumors were well established (in contrast with the experiment in Figure 6, in which treatment was started the day after tumor cell implantation). Mice in both large cohorts (without and with FcγR blockade) were randomly divided into 5 groups (n = 16 to 19 per group) for treatment with control mouse IgG, G6.31.2a, 4D5.2a, G6.31.2a plus 4D5.2a, or TG-VHS twice a week for 5 weeks (in contrast with the 6 weeks of treatment for the experiment in Figure 6). Figure 7A shows tumor development in each mouse in each treatment group from day 10 to day 25 after tumor cell implantation in mice without FcγR blockade (left panel) and with FcγR blockade (right panel). In both cohorts, TG-VHS produced the best tumor growth inhibition, and the combination treatment was better than 4D5.2a or G6.31.2a alone. Figure 7B compares the tumor sizes on day 25 among the different treatment groups in the same cohort and between the same treatment groups in the 2 cohorts. In the mice without FcγR blockade, tumor growth inhibition was significantly greater for TG-VHS treatment than for the combination treatment; by contrast, in the mice with FcγR blockade, the difference between TG-VHS and the combination treatment was notably smaller and statistically insignificant. In both cohorts, starting on day 18, mice in various treatment groups died or had to be euthanized owing to being moribund. Measurement of tumor growth was stopped after day 25, when more than 50% of the mice in the control antibody groups had died, leaving too few mice for detection of statistically significant effects; however, in both cohorts, all remaining mice in all treatment groups were continuously treated as scheduled until their death or completion of the planned treatment.

To confirm that 2.4G2 antibody treatment blocked the FcγRs on TAMs in the mice as expected, we analyzed tumor samples from mice euthanized on day 33, when a relatively large number of mice (n=30) happened to be identified as moribund and euthanized. Sixteen tumor samples from mice without FcγR blockade (1 treated with control IgG, 6 with G6.31.2a alone, 3 with 4D5.2a alone, 5 with the combination, and 1 with TG-VHS) were combined into one group, and 14 tumor samples from mice with FcγR blockade (2 treated with G6.31.2a alone, 2 with 4D5.2a alone, 4 with the combination, and 6 with TG-VHS) were combined into another group. We analyzed the percentage of total TAMs that had FcγR remaining available for interaction with a nonspecific goat-anti-rabbit IgG conjugated with PE fluorescence dye for detection by flow cytometry. As shown in Figure 7C, cancer cells (the cells with strong HER2 positivity) and TAMs (the cells with F4/80 positivity) represented 56.6±6.7% and 7.3±1.3%, respectively, of the total tumor cells harvested from the tumors in the cohort without 2.4G2 antibody treatment and 52.0±8.6% and 9.0±3.3%, respectively, of the total tumor cells harvested from the tumors in the cohort with 2.4G2 treatment. The differences between the 2 cohorts were statistically insignificant. However, the percentage of total TAMs with FcγR remaining available (FcγR-positive) for interaction with the PE-conjugated, nonspecific goat-anti-rabbit IgG was significantly lower in the tumor samples from the cohort with 2.4G2 antibody treatment (16.6±5.0%) than in the tumor samples from the cohort without 2.4G2 antibody treatment (32.7±6.2%). This finding corroborates that co-administration of 2.4G2 antibody blocked FcγRs on the TAMs in the mice and thereby diminished the access of the treatment antibodies to the FcγRs to induce ADCP.

Owing to the aggressiveness of the tumor model, all mice treated with control IgG died by day 40 (Figure 7D). On day 60, when all mice in the other treatment groups had died, 5 of the 18 mice (27.7%) in the TG-VHS group without FcγR blockade and 2 of the 19 mice (10.5%) in the TG-VHS group with FcγR blockade remained alive. Importantly, the median survival time for the mice treated with TG-VHS was 41 days without FcγR blockade compared with 28 days with FcγR blockade. This statistically significant 13-day improvement in median survival represents a 1.5-fold or 150% (41 days/28 days) improvement in median survival time. By contrast, for the other treatments, the median survival times were similar in the cohorts of mice with and without FcγR blockade: 25 days and 25 days, respectively, for control mouse IgG; 31.3 days and 28 days, respectively, for G6.31.2a; 26 days and 26 days, respectively, for 4D5.2a; and 35 days and 35 days, respectively, for the combination treatment. The fact that reduced median survival time in the mice with FcγR blockade was observed only for TG-VHS treatment supports our proposed working model of a role of VEGFA co-phagocytosis induced by the HER2-VEGFA BsAb via ADCP of HER2-overexpressing cancer cells in leading to an enhanced antitumor activity (Figures 3C and 5C).

## Discussion

In this work, we proposed a novel bispecific targeting strategy and assessed this strategy by developing and testing a pair of the BsAbs designed to induce co-phagocytosis of VEGFA (as an example of a soluble target in the TME) via inducing ADCP of cancer cells overexpressing HER2 (as an example of a solid target overexpressed on cancer cells). Our preclinical assessment of the strategy showed that our HER2-VEGFA BsAbs exerted greater anti-metastasis activity against aggressive tumor models, leading to a greater survival benefit, than did the parental anti-HER2 and anti-VEGFA antibodies used alone or in simple combination.

Depending on the desired functions or purposes, BsAbs can be engineered to be symmetrical or asymmetrical in molecular structures and can vary in molecular size from small scFv-based structures consisting of merely the variable regions of the parental antibodies linked by a flexible peptide to large complex IgG-like molecules retaining the IgG Fc domain of the parental antibodies (1, 3, 4, 34). In current work, our VHS BsAb platform was designed to include the IgG Fc domain, not only to ensure its interaction with FcγRs on macrophages for induction of ADCP but also to retain the pharmacokinetically favorable characteristics of antibody-based therapeutics (35, 36). It is worth mentioning that simultaneous bispecific binding to one soluble target in the TME and one solid target on the cancer cell surface is critical for our proposed mechanism of action. However, some BsAb platforms, despite their strong binding to 2 different targets individually, have a limited capacity in binding to the 2 different targets simultaneously. This can happen when respective binding sites on the hypervariable complementarity-determining regions within the variable regions of the BsAb are in proximity leading to intramolecular steric hindrance between the 2 targets (37). Within our VHS platform, we observed remarkably less competition between HER2 and VEGFA for simultaneous binding to the BsAb than was previously reported for a “2-in-1” BsAb design (37). One potential disadvantage of the VHS platform, however, is its relatively large size as a result of scFv fusion, which may theoretically impact the penetration of the BsAb into solid tumors. Nevertheless, the size does not appear to be a major concern given that our BsAbs induced strong tumor responses in our in vivo studies.

Using a cell co-culture system, we found that VEGFA-EGFP signal was detected both in cancer cells (4T1-HLC cells) and in macrophages (RAW264.7) upon TG-VHS treatment, which supports our proposed working model (Figure 3C) in which VEGFA co-phagocytosis upon TG-VHS treatment occurs via a 2-step process. In step 1, soluble VEGFA in the medium was anchored onto cancer cells following bispecific binding of TG-VHS to VEGFA and HER2. In step 2, the anchored VEGFA was phagocytized along with the targeted cancer cells by RAW264.7 macrophages via ADCP induced by TG-VHS. When we expanded the antibody treatment to overnight and used primary BMDM cells that exhibit a higher phagocytic activity than the established RAW264.7 macrophages, we further confirmed a higher level of VEGFA uptake by the BMDM cells in co-culture with cancer cells after treatment with the BsAb than with the combination of anti-VEGFA and anti-HER2 antibodies. Interestingly, we also found that the polarized BMDM cells (M1 type or M2 type) mediated a higher level of TG-VHS-induced VEGFA co-phagocytosis than did the unpolarized BMDM cells that are considered resting macrophages in an inactive state or M0 type. This finding is consistent with the observation of higher levels of FcγR in the polarized macrophages than in the unpolarized BMDM cells. This finding is important because the TAMs in the TME often have been polarized, with most of them being of the M2 type macrophages. Of note, although the 2.4G2 antibody can recognize both FcγRIII/CD16 and FcγRII/CD32 (27, 28), the increase in the FcγRs in the polarized BMDM cells were reported to be mainly FcγRII/CD32 (38).

In our studies, we developed 2 versions of HER2-VEGFA BsAbs—TB-VHS, a fully humanized IgG1 antibody, and TG-VHS, a murine IgG2a antibody counterpart that functions analogously to human IgG1 in terms of the IgG effector functions in the species of the host (mouse) (39). Our findings from the nude mice with 4T1-HL tumors provided evidence that TB-VHS worked better than trastuzumab and bevacizumab alone or in combination through mechanism(s) without participation of the host immune system. Our findings from the immunocompetent hmHER2Tg mice with D5-HER2 tumors further showed a clear survival benefit of TG-VHS over the combination treatment. It is worth discussing that when we assessed therapeutic responses to the antibody treatments in mice that started with a well-established aggressive tumor (Figure 7), overall survival in the treatment groups, except in the control group, was lower than in the corresponding treatment groups started early after tumor cell implantation (Figure 6). Owing to this lower overall survival, the improvement in overall survival between the TG-VHS-treated mice with and without FcγR blockade seemed small when the experiment was ended on day 60. Nevertheless, the median survival time, which is often used in clinical trials to evaluate the effectiveness of a new treatment, was shortened from 41 days in the mice treated with BsAb alone to 28 days in the mice treated with BsAb plus FcγR blockade. This statistically significant 13-day difference represents a close to 150% or 1.5x difference in median survival time and can be converted approximately to 533 days in human (one mouse day equivalent to around 41 human days (40). The median survival times were not shortened in the other treatment groups plus FcγR blockade. This finding supports a functional role of FcγR in mediating an enhanced activity of TG-VHS compared with other treatments. It is acknowledged that the 2.4G2 antibody might also block CD16 on NK cells, leading to inhibition of antibody-dependent cell-mediated cytotoxicity (i.e., ADCC), which would affect tumor response not only to TG-VHS but also to 4D5.2a alone or in combination with G6.31.2a; however, no significant difference was found in the antitumor activity of 4D5.2a alone or G6.31.2a plus 4D5.2a between the cohorts receiving and not receiving 2.4G2 antibody. Together, the findings support the conclusion that VEGFA co-phagocytosis via ADCP is a key mechanism by which TG-VHS exerts strong antitumor activity.

VEGFA is a key modulator of host innate immune response with suppressive effects in addition to its pro-angiogenic activities (16, 17). Several mechanisms have been reported by which VEGFA suppresses tumor immune responses, including promoting M2 polarization of macrophages to secret anti-inflammatory cytokines, such as IL-10 and TGF-β (41), inhibiting dendritic cell maturation (42), upregulating immune checkpoint regulators, such as IDO (42) and PD-L1/2 in myeloid cells (42, 43), and recruiting myeloid-derived suppressor cells and regulatory T cells into tumors (44, 45). Anti-VEGFA therapy can also improve host adaptive tumor immune response via vascular normalization, leading to increased tumor-infiltrating lymphocytes, such as CD4^+^ and CD8^+^ T cells (43, 46). We found that the BMDM cells express VEGFRs in our studies, consistent with the findings in literature (41, 47). Strong inhibition of VEGFA by TG-VHS may have modulated these activities and thereby promoted an anti-tumor immune response in the hmHER2Tg mice. It is of interest to note the rejection of rechallenge with the parental D5 tumor cells (without HER2) in those mice that were inoculated with D5-HER2 tumor cells and became tumor-free after TG-VHS treatment. However, D5 tumor cells express a low level of MHC-I (H-2K^b^ and H-2D^b^) and D5-HER2 tumor cells express even a lower level of that than parental D5 cells (31, 32). This finding, therefore, should be interpreted with caution regarding whether the presence of effector CD8^+^ T cells contributed to tumor-free survival of the mice after TG-VHS treatment. Nevertheless, the finding that effector CD8^+^ T cell population was increased upon TG-VHS treatment corroborates that inhibition of VEGF-A can enhance host immune responses by modulating immune cells, including T cell function.

In summary, our findings support the concept of a novel mechanism of action exerted by the HER2-VEGFA BsAbs in VHS platform that leads to a survival benefit. Given that the BsAbs improved the median survival in mice with aggressive murine tumors that are lethal due to massive metastases, it is reasonable to expect that the humanized TB-VHS might produce similar outcomes in patients with late-stage metastatic cancer. Our results warrant clinical testing of the BsAb strategy for patients with HER2-overexpressing cancers across a variety of cancer types, in particular cancers that respond to bevacizumab, such as colorectal cancer. Furthermore, our proof of the concept of a novel mechanism of action exerted by HER2-VEGFA BsAbs suggests that the VHS platform could be applied to develop additional BsAbs with one target overexpressed on the cancer cell surface (e.g., EGFR, CD19 or CD20) and the other soluble target abundant in the TME (e.g., IL-10 or TGFβ) to achieve desired activities against various types of tumors.

## Methods

### Sex as a biological variable

Both male and female mice were included in this study and findings were similar in both sexes.

### Animals

Female Swiss nude mice were ordered from a mouse colony facility maintained by the Department of Experimental Radiation Oncology, MD Anderson Cancer Center. The original transgenic breeding pairs of hmHER2Tg mice (C57BL/6 background) were provided by Dr. Louis M. Weiner (Georgetown University Medical Center) (31, 32). The hmHER2Tg mice have been maintained at MD Anderson Cancer Center by the principal investigator’s team since then. Both male and female hmHER2Tg mice were used in this study.

### Mouse tumor models

Two implantable mouse tumor models were used in this study. The 4T1-HLC tumor cells and the co-implanted NIH3T3-V cells were generated via lentiviral transduction of the respective parental cells with the respective cDNA construct of interest (HER2, firefly luciferase, EGFP, and VEGFA-EGFP fusion protein) packed using the pLEX lentivirus packing system. 4T1-HL cells (1×10^5^ cells) were implanted along with NIH3T3-V cells (5×10^5^ cells) into the mammary fat pad of Swiss nude mice (4-6 weeks old). D5 tumor cells and D5 tumor cells expressing HER2 (D5-HER2) were provided by Dr. Louis M. Weiner (Georgetown University Medical Center) (31, 32). D5 cells and D5-HER2 cells (2.5×10^5^ cells/mouse) were implanted subcutaneously in the mice (8-12 weeks old). Tumor development and tumor response to various treatments were monitored twice a week.

### Mouse treatments and analysis of tumor development and mouse survival in response to the treatments

TB-VHS, TG-VHS, 4D5.2a, and G6.31.2a antibodies were produced in house. The details are described below under “Engineering and production of BsAbs with the VHS platform and respective parental antibodies”. Trastuzumab and bevacizumab were purchased from the pharmacy at MD Anderson Cancer Center. Non-pharmaceutical grade 2.4G2 anti-mouse CD16/CD32 antibody was purchased from Selleckchem.

Female nude mice were used for assessing the therapeutic activity of TB-VHS and respective parental antibodies with details described in relevant figure legends. Tumor development and tumor response to various treatments were monitored weekly in live mice by IVIS imaging. Both male and female hmHER2Tg mice were used for assessing the therapeutic activity of TG-VHS and respective parental antibodies with details described in relevant figure legends. Tumor volumes were measured twice a week with calipers and calculated using the formula volume = π/6 × ab^2^ (a: length; b: width, a > b). Mice were euthanized when the animals became moribund or when the tumor size in the mice reached greater than 2.0 cm in diameter.

### Tumor rechallenge experiment

Age-matched treatment-naïve mice and the mice cured after various treatments in the experiment underwent subcutaneous implantation of equal numbers of parental D5 tumor cells (2.5×10^5^ cells/mouse); this was followed by close observation of tumor development and tumor size measurement in the mice as described above.

### Immunophenotypic analysis of mouse spleen specimens

Mouse spleen specimens were minced mechanically and then passed through a 70-μm mesh cell strainer to isolate single cells. Single cell suspensions (0.5-1×10^6^ cells/sample) were first subjected to FcγR blockade with TruStain FcX and co-stained with Ghost Dye Violet 510 (Tonbo Biosciences) and then were stained with respective fluorescently conjugated antibodies (1-2 μL in 100 μL of flow cytometry staining buffer [0.5% BSA in PBS]). The fluorescently conjugated primary antibodies included PE-conjugated anti-mouse CD8a (clone 53-6.7), violetFluor 450-conjugated anti-human/mouse CD44 (clone IM7), PE-Cy7-conjugated anti-mouse CD3e (clone 145-2C11), APC-conjugated anti-F4/80 (clone RM8), and PerCP/Cy5.5-conjugated anti-CCR7, which were purchased from Tonbo Biosciences or BioLegend. After staining for 30 min at 4°C in the dark, the cells were washed twice with cytometry staining buffer and then analyzed by using a Becton Dickinson Flow Cytometer. The flow cytometry data were analyzed by using FlowJo software (v10) to select subpopulations of cells gated successively on the fluorescence of interest.

### Analysis of pharmacological blockade of FcγR ex vivo

Tumor specimens collected from the cohorts of mice treated with or without blockade of FcγR (2.4G2 antibody, 100 µg/mouse) were processed for single cell suspension and then stained for flow cytometry as described above with fluorescently conjugated primary antibodies, including BV421-conjugated anti-HER2 antibody (BioLegend), APC-conjugated anti-F4/80 antibody (BioLegend), and PE-conjugated nonspecific goat-anti-rabbit IgG antibody (Santa Cruz Biotechnology).

### Harvest of BMDM cells from mice

Primary BMDM cells were harvested from normal mice following protocols in literature (48, 49).

### Cell culture

All cell lines used in the study were obtained from the American Type Culture Collection (ATCC) unless otherwise stated. All cell lines except primary BMDM cells were cultured in DMEM/F12 medium (50/50, v/v) supplemented with 10% FBS, 2 mM glutamine, 100 U/mL penicillin, and 100 μg/mL streptomycin. BMDM cells were cultured in RPMI 1640 medium supplemented with 50 ng/mL M-CSF (SinoBiological), 10% FBS, 2 mM glutamine, 100 U/mL penicillin, and 100 μg/mL streptomycin. All cells were incubated at 37°C with 5% CO2 in a humidified atmosphere.

### Polarization of BMDM cells in culture

For inducing M1 type polarization, BMDM cells were treated with a combination of IFNγ (50 ng/mL, SinoBiological) and LPS (1:1000 from the 500x stock, eBioscience) for 12 hours in culture. For inducing M2 type polarization, BMDM cells were treated with a combination of IL-4, IL-10 and IL-13 (SinoBiological) at 50 ng/mL of each for 24 hours in culture. The outcome of polarization was confirmed by flow cytometry analysis as described above after staining with anti-CD86 antibodies (BioLegend) for M1 type macrophages or anti-CD206 antibody (BioLegend) for M2 type macrophages, respectively.

### Analysis of BsAb-induced co-phagocytosis of VEGFA via ADCP of HER2-overexpressing cancer cells in co-culture with macrophages

For induction of co-phagocytosis of VEGFA-EGFP by TG-VHS in co-culture of 4T1-HLC cells with RAW264.7 mouse macrophages (at a 1:5 ratio of cell number), equal volumes of conditioned medium collected from culture of NIH3T3-VG fibroblasts or from culture of NIH3T3-G fibroblasts were added respectively to the co-culture of 4T1-HLC and RAW264.7 cells on a 6-well culture plate supplemented with fresh medium. The co-cultured cells were then treated with control mouse IgG, G6.31.2a, 4D5.2a, G6.31.2a plus 4D5.2a, or TG-VHS for 2 hours in a 37°C incubator with 5% CO_2_. The culture medium was then removed, and the cells were washed and harvested. After additional FcγR blockade with TruStain FcX antibody (anti-mouse CD16/CD32) (BioLegend), the cell samples were stained with APC-conjugated anti-F4/80 antibody for flow cytometry analysis as described above.

For induction of co-phagocytosis of pHrodo green-labeled recombinant VEGFA by TG-VHS in co-culture of cancer cells with primary BMDM cells, recombinant VEGFA (50 µg, Sinobiological) was labeled with pHrodo green fluorescent dye (100 µg) according to the protocol of pHrodo iFL microscale labeling kit provided by the vendor (Invitrogen). The labeled VEGFA (0.5 µg/10 µL) was added to BMDM cells (2×10^5^ cells) in 500 μL medium cultured without or with D5-HER2 tumor cells (4×10^5^ cells, i.e., at a 1:2 ratio of cell number) in the presence of mouse IgG, G6.31.2a plus 4D5.2a or TG-VHS (at 5µg/mL of each) in a FACS tube in a 37°C incubator overnight. The cell samples were then subjected to FcγR blockade and staining with APC-conjugated anti-F4/80 antibody for flow cytometry analysis as described above.

### Laser scanning confocal microscopy

Laser scanning confocal microscopy was performed on a Zeiss LSM880 using the Airyscan detector and 40X 1.2 N.A. water apochromat objective with a 2X zoom. Images were captured as Z-stacks with an interval of 1 µm, 12 µm range and a pixel size of 80 nm. Airyscan processing was performed in Zen Black using default strength parameters. Z-stacks were processed in Zen Blue image analysis software to generate maximum intensity projections as indicated. Experimental and control samples were imaged and processed in parallel using the same acquisition (e.g., transmission, gain, etc.) and processing parameters.

### Engineering and production of BsAbs with the VHS platform and respective parental antibodies

The complementarity-determining region sequences used for construction of TB-VHS, TG-VHS, 4D5.2a, and G6.31.2a expression vectors were obtained from the public domain and are published (50). The VH and VL sequences of 4D5.2a were based on those of trastuzumab, and the VH and VL sequences of G6.31.2a were based on those of G6.31, but the Fc domain of 4D5.2a and G6.31.2a was engineered in a mouse IgG2a framework to ensure comparability with the Fc domain of TG-VHS for IgG effector functions. Codon-optimized DNA fragments encoding the antibody genes were synthesized, constructed, subcloned into a homemade construct, and expressed in CHO-DG44 cells, which were cultured in chemically defined serum-free medium. The antibodies produced by the CHO-DG44 cells were then purified by subjecting the conditioned medium to binding to protein A–ceramic HyperD F affinity chromatography sorbent (Pall Laboratory), followed by acid elution with 0.1M glycine (pH 2.6) and immediate neutralization with 1M Tris-HCl (pH 11). Samples of protein A–purified antibodies were quantified using the Pierce Coomassie Plus (Bradford) colorimetric protein assay (Thermo Fisher Scientific) and were separated by SDS-PAGE, followed by gel staining with Coomassie blue and then soaking in a de-staining solution overnight to clean background. The purified antibodies were dialyzed with PBS, then sterilized via filtration through a 0.2-μm membrane device, and kept at -80°C in aliquots before use.

### Characterization of BsAbs for binding to HER2 and VEGFA individually and competitively

Binding of purified TB-VHS to its respective targets (HER2 and VEGFA) was measured in a cell-free PBS solution by ELISA. For detecting HER2 binding, 96-well microplates coated with HER2 ECD recombinant protein (SinoBiological) were used to capture TB-VHS or the parental anti-HER2 antibody. The captured antibodies were then detected by HRP-labeled anti-human IgG antibody (Jackson ImmunoResearch) in an ELISA. For detecting VEGFA binding, 96-well microplates coated with rabbit anti-human or anti-mouse IgG Fc antibody (Jackson ImmunoResearch) were used to capture TB-VHS or the parental anti-VEGFA antibody. The captured antibodies were then incubated with biotinylated human or mouse VEGFA (G&P Biosciences) and detected by streptavidin-HRP (Invitrogen) in an ELISA.

For assessing competitive binding of the 2 targets to TB-VHS, either TB-VHS (5 nM) was incubated in a solution with biotinylated VEGFA at a fixed concentration (5 nM) and HER2 ECD recombinant protein at increasing concentrations at 4°C for 1 hour, or TB-VHS (5 nM) was incubated in a solution with HER2 ECD recombinant protein at a fixed concentration (5 nM) and biotinylated VEGFA (mixtures with unlabeled VEGFA) at increasing concentrations at 4°C for 1 hour. The reaction products were then applied to separate 96-well microplates coated with anti-human IgG Fc antibody (to capture TB-VHS). VEGFA bound to TB-VHS was detected by streptavidin-HRP in an ELISA, whereas HER2 ECD bound to TB-VHS was detected by streptavidin-HRP after incubation with a biotinylated anti-HER2 antibody (Invitrogen).

The capacity of TB-VHS or TG-VHS for binding to VEGFA following binding of the BsAbs to HER2 overexpressed on live cells was measured by flow cytometry. After incubation at 4°C for 1 hour with TB-VHS or TG-VHS or with the respective parental antibodies, HER2-overexpressing SKBR3 human breast cancer cells were incubated with FITC-labeled anti-human IgG antibody or FITC-labeled anti-mouse IgG antibody (Jackson ImmunoResearch); FITC-avidin (R&D) plus biotinylated human VEGFA; or FITC-avidin plus biotinylated mouse VEGFA. The cells were then subjected to flow cytometry analysis as described above.

### Statistics

Statistical analyses were performed with GraphPad Prism 10 software. The log-rank test was used to compare the survival distributions of two or more groups of mice after various treatments. Experimental data in the animal studies are presented with the mean values with standard error. A two-tailed unpaired Student’s t test was used to compare 2 groups of tumor volumes in mice and the ex vivo and in vitro samples after various treatments. Fisher’s exact test was used to compare the percentage of mCherry (or EGFP)-positive or -negative cells in macrophages (F4/80-positive cells).

### Study approval

All mouse experiments were performed in accordance with guidelines and protocols approved by the Institutional Animal Care and Use Committee of The University of Texas MD Anderson Cancer Center.

## Data availability

All data generated and analyzed during this study are available in the Supporting Data Values file. Materials can be obtained upon reasonable request by contacting the corresponding author.

## Author contributions

ZF conceived the original idea, acquired the funding, supervised the project, and wrote the manuscript.

YL, SQ, and ZF designed the study.

YL and SQ performed the experiments.

YL, SQ, and ZF analyzed the data.

All authors approved the final version of the manuscript.

YL was given first listing, as YL initiated the project.

## Acknowledgments

This study was supported in part by NIH grant R01CA262288 and a grant from the Breast Cancer Research Foundation (BCRF-18-051, 19-051, 20-051, 21-051, 22-051, 23-051, and 24-051). The work was also supported in part by the NIH through MD Anderson’s Cancer Center Support Grant, CA016672, which supported the animal, FACS and confocal cell imaging studies performed in this study. We are grateful to Dr. Louis M. Weiner of Georgetown University Medical Center for providing the hmHER2Tg mouse model and the cell models (D5 and D5-HER2). We would like to thank Dr. Collene Jeter of the Advanced Microscopy Core at MD Anderson Cancer Center for assistance with confocal cell imaging, Dr. Mahrima Parvin at Dr. Zhen Fan’s lab for assistance with BMDM collection, and Stephanie Deming of the Research Medical Library, MD Anderson Cancer Center, for editing this manuscript.

---

**Supplemental Figure 1.**
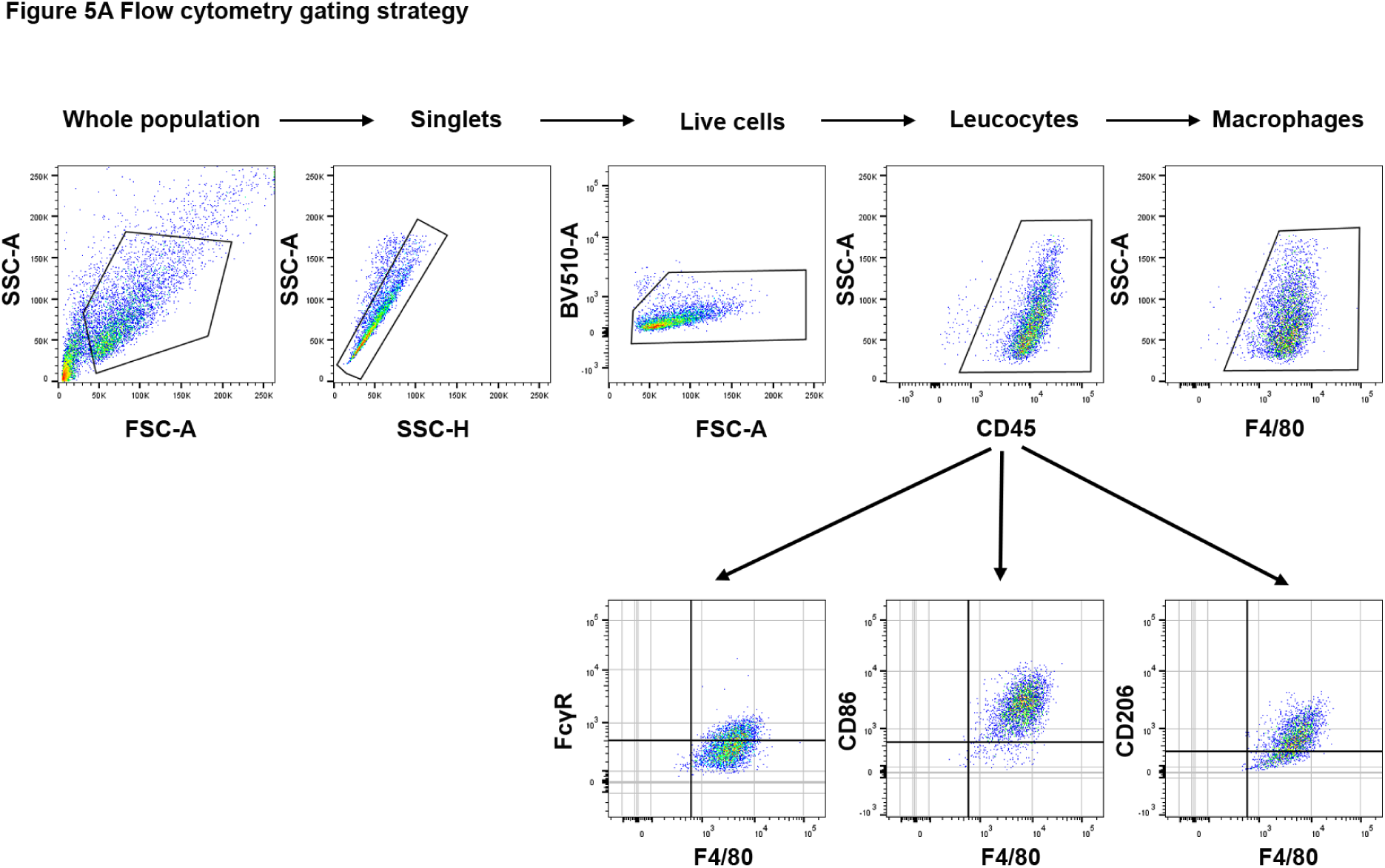

**Supplemental Figure 2.**
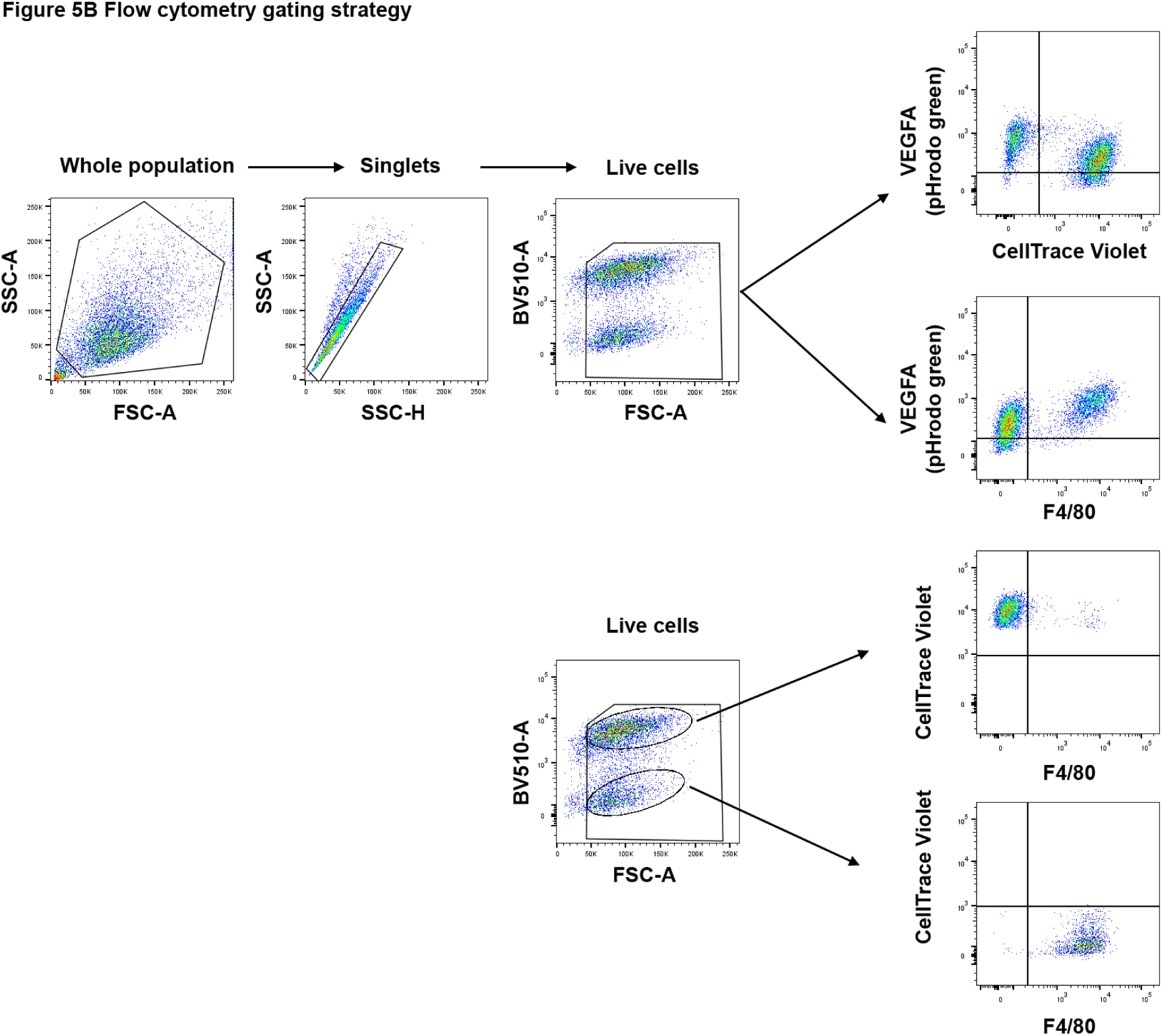

